# Mating enhances immune function of *Drosophila melanogaster* populations against bacterial pathogens

**DOI:** 10.1101/823211

**Authors:** Nitin Bansal, Biswajit Shit, Aparajita, Tejashwini Hegde, Rochishnu Dutta, Nagaraj Guru Prasad

**Author notes:** Corresponding author (RD). **Author Contribution:** N.B., A. and N.G.P. conceived and designed research; N.B., B.S., T.H., and A. generated flies and executed immunity assays; N.B. and R.D. analyzed data and wrote initial draft; N.G.P. corrected draft manuscript. **Data Archive:** Data has been added as a supplementary material. There is no other data to be archived. **Ethics statement:** The study involved laboratory reared populations of *Drosophila* that does not fall under the IUCN Red List of Threatened Species or Schedule Species Database of Government of India and hence is exempted from any approval of National Biodiversity Authority (Government of India).

## Abstract

Immunity and reproduction are two important processes that affect fitness of an organisms. Sexual activity has been previously shown to determine the degree to which a population is able to survive various infections. While many studies have demonstrated a trade-off between immune function and reproduction, many studies have found synergistic relation between the two fitness determinants. Besides it is generally hypothesised that sexes may differ in immunity due to relative cost they incur during reproduction with males losing in immunity to rather increase their reproductive success. In this study, we test the effect of immune function on the survival of mated and virgin replicates of a large outbred baseline *D. melanogaster* population that was infected with four different bacterial infections. We find enhanced survival in mated flies than virgin flies in response to all four bacterial infections across all replicates. While investigating sexual dimorphism in immune function, we find no difference in sexes in their ability to survive the imposed bacterial infections. Synergistic interaction between reproduction and immunity may exist if it improves Darwinian fitness of either sexes of a population under selection, and are not necessarily limited by each other due to trade-off over finite resources.

## Introduction

Post mating immune function is the ability of an organism to resist harmful pathogens after becoming susceptible to infection due to mating (Oku et al., 2019). The probability to survive possible infection post mating, is not always correlated to sub-organismal immune response (measured in terms of encapsulation, haemocyte load, prophenoloxidase and phenoloxidase activity, lytic activity, gene expression, etc.) to mating process itself (Adamo, 2004; Fedorka et al., 2007; Oku et al., 2019; Zuk and Stoehr, 2002). We, therefore, only deal with post mating immune function measured in terms of survival (probability of surviving) or hazard function (probability of death) as a measure of overall immunity in this study. Although mating is generally believed to trade-off with post mating immune function as a result of conditional partitioning of finite resources (Adamo, 2004; Schwenke et al., 2016; Zuk and Stoehr, 2002), mating is often found to enhance immune functions in many invertebrates (Barribeau and Schmid-Hempel, 2017; Msaad Guerfali et al., 2018; Shoemaker et al., 2006; Valtonen et al., 2010; Worthington and Kelly, 2016). However, studies of post mating immune function in *Drosophila melanogaster* have thrown up mixed results in the past, especially varying with bacteria used to infect the fruit flies, nature of sexual experience of the fruit flies and the time of infection post mating.

Mated *D. melanogaster* male flies have been found to lose resistance to non-pathogenic *Escherichia coli* with increase in sexual activity in one study (McKean and Nunney, 2001) while in another study (Wigby et al., 2008), effectiveness of the immune function triggered by *Escherichia coli* infection did not vary between mated and virgin *D. melanogaster* females. On the contrary, in three *D. melanogaster* populations, mated *D. melanogaster* males were found to be more resistant to highly virulent *Pseudomonas entomophila* infection than virgin males while mated and virgin males survived the less virulent *Staphylococcus succinus* infection equally well (Gupta et al., 2013). Mated *D. melanogaster* females were found to suffer higher mortality than virgin females due to pathogens such as *Providencia rettgeri* and *Providencia alcalifaciens* while no difference between mated and virgins were observed in infections from *Pseudomonas aeruginosa* and *Enterococcus faecalis* (Short and Lazzaro, 2010). Effect of *Pseudomonas aeruginosa* infection on mated *D. melanogaster* females, however, was found to be detrimental in another study (Fedorka et al., 2007). These studies have several limitations such as (a) using one or two types of bacteria that differ in virulence leading to variability in results, (b) pathogens that are not natural to hosts (c) using either male or female fruit flies thereby making any generalisation sex specific (d) one study with four bacteria (Short and Lazzaro, 2010) used multiple isofemale *D. melanogaster* lines that lack within-line natural genetic variation. We are not sure whether the effect of inbreeding over many generations could possibly confound the isofemale flies’ response to infection (problem discussed in Adamo, 2004; McKean and Nunney, 2008; Rigby et al., 2002). Therefore, to study the effect of mating on post mating immune function in *D. melanogaster*, we (a) used multiple bacteria that are natural pathogens to fruit flies (b) used both Gram-positive and Gram-negative bacteria (c) used both male and female fruit flies (d) infected flies from a large outbred *D. melanogaster* wildtype population that has been maintained in laboratory under no particular selection pressure.

Studies investigating immunity-reproduction trade-offs are predominantly sex specific due to the assumption that sexes may commit their resources to immunity differently during reproduction (McKean and Nunney, 2001; Rolff, 2002). Theory suggests that males will spend more resources in mating at the expense of immunity, making them more vulnerable to infections (Zuk and Stoehr, 2002). Meta-analysis on sex difference in invertebrate immunity, however, found no sexual dimorphism (Kelly et al., 2018; Letitia et al., 2000). While differences in post mating immunity between sexes has been observed in some cases, any trend generalising a sex biased immunocompetence remains elusive (Kelly et al., 2018; McKean and Nunney, 2005). In this study, we also test whether mated *D. melanogaster* females differ from mated males in their post mating immune function.

We challenged mated and virgin fruit flies of both sexes with sublethal doses of two Gram-positive and two Gram-negative bacteria. In case the mated flies experience more mortality than the virgin flies, it would suggest that the mated flies are more susceptible to infections and vice versa. The result will then provide us whether reproduction and immune function have antagonistic or synergistic relationship. Any observed difference in survival between males and females of the fruit fly population will help us determine whether sexual dimorphism exist in post mating immune function. Our objectives for this study were as follows:

1. To investigate survival of mated and virgin wild type *D. melanogaster* flies facing bacterial infections.
2. To investigate whether sexual dimorphism exists in immunological performance of the *D. melanogaster* flies after infection.

## Methods

### *D. melanogaster* population

We used a large (N = 2800) outbred laboratory-adapted BRB (Blue Ridge Baseline) (Singh et al., 2015) population of *D. melanogaster* (henceforth, BRB) that has been maintained in five replicates in our laboratory. We used three out of the five BRB replicates, each of which were 188 generations old independent lineage at the time of our experiment but as a whole represents a genetically variable population. These distinct replicates having common ancestors are good fit for the present study since their response to bacterial infection will not be constrained by any differences in ancestral conditions or long-term selection pressure.

For our experiment, we collected eggs at 70 eggs per vial (90-mm length × 25-mm diameter) in 8-10 ml (Gupta et al., 2013) of banana-jaggery-yeast medium from all three replicates of the stock population (for more details, see Singh et al. (2015)). We chose some vials randomly for virgin collection. Virgins were gathered every 6 hours on days 10^th^ and 11^th^ post egg collection and the sexes were housed separately in single sex vials at a density of 10 per vial. To obtain mated flies, we kept the remaining vials for flies to eclose, get sexually active and mate within their rearing vials till the 12^th^ day post egg collection. Here, we assume that the sexually active flies will mate multiple times during two days (10^th^−12^th^) period post ecclosion (since flies start mating 8 hours post eclosion (Gupta et al., 2013)).

### Bacterial populations and infection protocol

We used four bacteria, two Gram-positive bacteria (*Enterococcus faecalis (Ef)* and *Staphylococcus succinus (Ss))* and two Gram-negative bacteria (*Pseudomonas entomophila (Pe)* and *Providencia rettgeri (Pr))* in this study. *Ef* and *Pr* bacteria were provided by Prof. B Lazzaro (Juneja and Lazzaro, 2009; Lazzaro et al., 2006), *Pe* was provided by Prof. P. Cornelis (Vodovar et al., 2005) and *Ss* was isolated from wild caught *Drosophila melanogaster* in our laboratory (Singh et al., 2016). Primary bacterial culture of each bacteria used in this study was derived from the respective bacterial glycerol stocks and setup in sterilised falcon tubes containing 5 ml of Luria Bertini (LB) broth medium. These falcon tubes were kept at bacteria specific optimum growth temperature (see Supplementary Information) overnight for bacteria specific growth period (see Supplementary Information) at 150 revolutions per minute in bacterial incubator (Innova 42 Incubator Shaker, New Brunswick Scientific), a day before the infection assay. From primary culture, 200 ul of solution was resuspended in fresh 10 ml of LB to make secondary culture. The secondary culture was subsequently pelleted down using a centrifuge (Centrifuge 5430R, Eppendorf) at 7200 revolutions per minute for around 10 minutes and finally suspended in desired volume of 10 mM MgSO_4_ solution to obtain a final optical density of 1 at 600 nm. After lightly anesthetizing the flies using CO_2_, each fly was infected by pricking the lateral side of the thorax with a stainless steel Minutien pin (Fine Science Tools, CA) of 0.1 mm diameter dipped in bacterial suspension in 10 mM MgSO_4_. We imparted the sham infection (control) using Minutien pin dipped only in sterile 10 mM MgSO_4_ following Gupta et al. (2013) protocol.

### Survivorship assay

At the end of 12^th^ day post egg collection, we infected our experimental flies with different bacteria and sham treatments within 3 hours. The sample size for the experiment was 6000 [100 × 2 (sex) × 2 (mating status) × 3 (replicates) × 5 (4 bacterial + 1 sham treatment)]. After infections, flies were separately housed in different Plexiglass cages provided with food plates (each containing 100 individuals) according to sex, treatment type and replicate identity. Mortality was monitored for at least 96 hours post infection.

### Statistical analysis

For statistical analysis we recorded the variables (1) sex, (2) replicate affiliation, (3) mating status, (4) bacterial infection type and (5) time till death of the BRB flies. We combined two of the variables, (3) mating status and (4) bacterial infection type, to form 10 treatments (namely, ‘Mated *Ef*’’, ‘Virgin *Ef*’’, ‘Mated Sham’, ‘Virgin Sham’, etc.) that were subsequently used in analysis. We censored the mortality data by assigning value of 1 (observed death) or 0 (no death within observation period). We used Cox proportional hazard model to examine simultaneous effect of all predictor variables on the hazard rate of BRB flies. For univariate survival analysis and plotting survivorship curves, we used Kaplan-Meier method. We performed both Cox proportional hazard analysis and Kaplan-Meier analyses using ‘Survival’ package (Therneau, 2015) in R (Team, 2019). We used open source ‘ggkm’ package to plot survivorship curves in R.

## Results

Cox proportional hazard test involved 3 predictor variables: treatment, replicate, and sex (details in Supplementary Information). In this multivariate analysis, risk of death for ‘Virgin Sham’ treatment and ‘Mated Sham’ treatment was negligible and not significantly different from each other. However, infected mated and virgin BRB flies had significantly higher risk of mortality when compared with ‘Virgin Sham’ treatment. We also found that the female flies were no different to male flies in surviving the bacterial infections (see Supplementary Information). In the same analysis, we found that the second BRB replicate had relatively lower risk of death due to bacterial infection (hazard ratio = 0.84, p = 0.0007) compared to the first replicate while the third replicate faced similar hazard as the first replicate. Such difference is not unexpected since each BRB replicate has undergone long independent evolution in laboratory cultures. However, since the results of each BRB replicate in Kaplan-Meier survival analysis were consistent (see Supplementary Information), we pooled all three BRB replicate data to present a single survivorship curve (see Figure 1).

**Figure 1.**
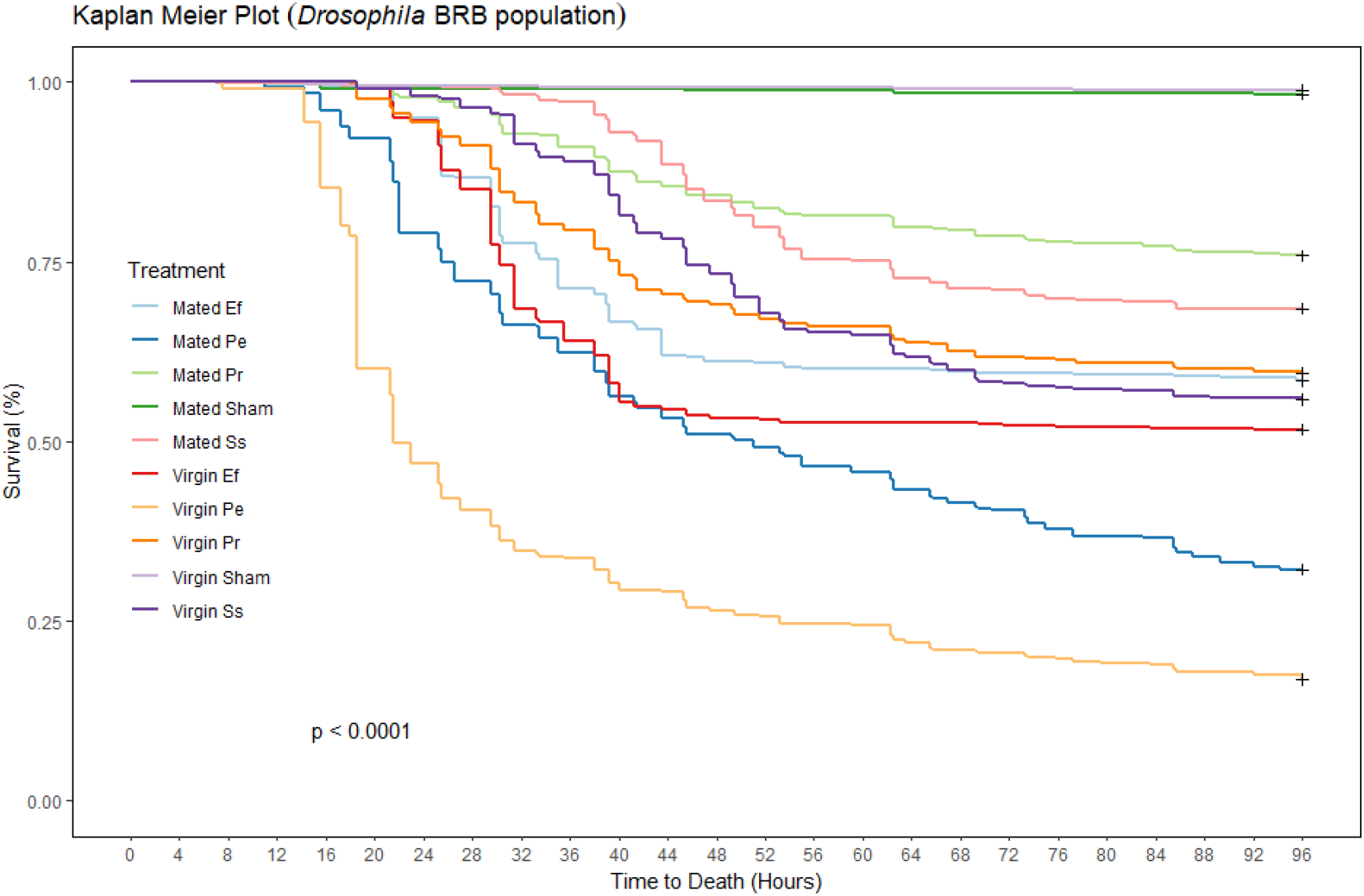
Kaplan-Meier plot showing that mated *D. melanogaster* BRB flies are significantly better than virgin flies at surviving infection from two Gram-negative bacteria, *Pseudomonas entomophila* (*Pe*) and *Providencia rettgeri* (*Pr*) and two Gram-positive bacteria, *Enterococcus faecalis* (*Ef*) and *Staphylococcus succinus* (*Ss*).

Kaplan-Meier survival analysis suggests that mated BRB flies survived significantly better than virgin flies infected with sub-lethal doses of *Enterococcus faecalis*, *Pseudomonas entomophila*, *Providencia rettgeri* and *Staphylococcus succinus* infections (see Figure 1). This mated-outperforming-virgin trend was consistent even when we investigated BRB flies’ survivorship for each bacterial infection separately (see Table 1 and Supplementary Information). Both the sham treatments (for mated and virgin control) had negligible mortality in this study indicating any mortality in other treatments were only due to bacterial infections (see Figure 1). *Pseudomonas entomophila* turned out to be the most virulent among infecting bacteria (see Table 1) for BRB flies with median time to death at 21.5 hours for virgin BRB flies and 51 hours for mated BRB flies. Kaplan-Meier survival analysis using sex as predictor variable showed that both sexes were equally adept at withstanding bacterial infections (see Figure 2). For individual bacterial infections analysed separately, female and male survival were not significantly different (see Supplementary Information).

**Figure 2.**
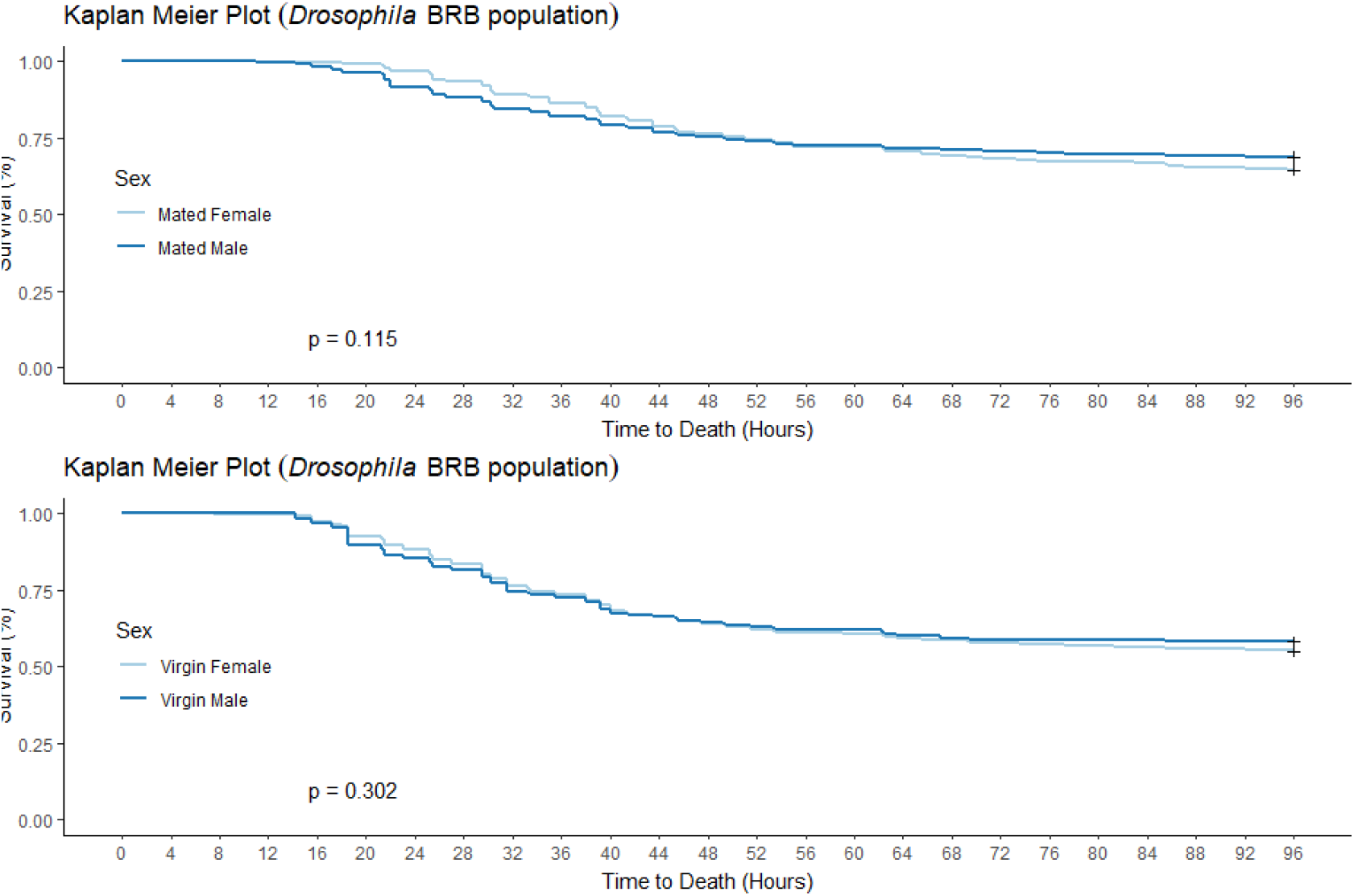
Kaplan-Meier plot showing no differences in male and female survivorship when *Drosophila* BRB flies are infected with four bacteria.

**Table 1.**
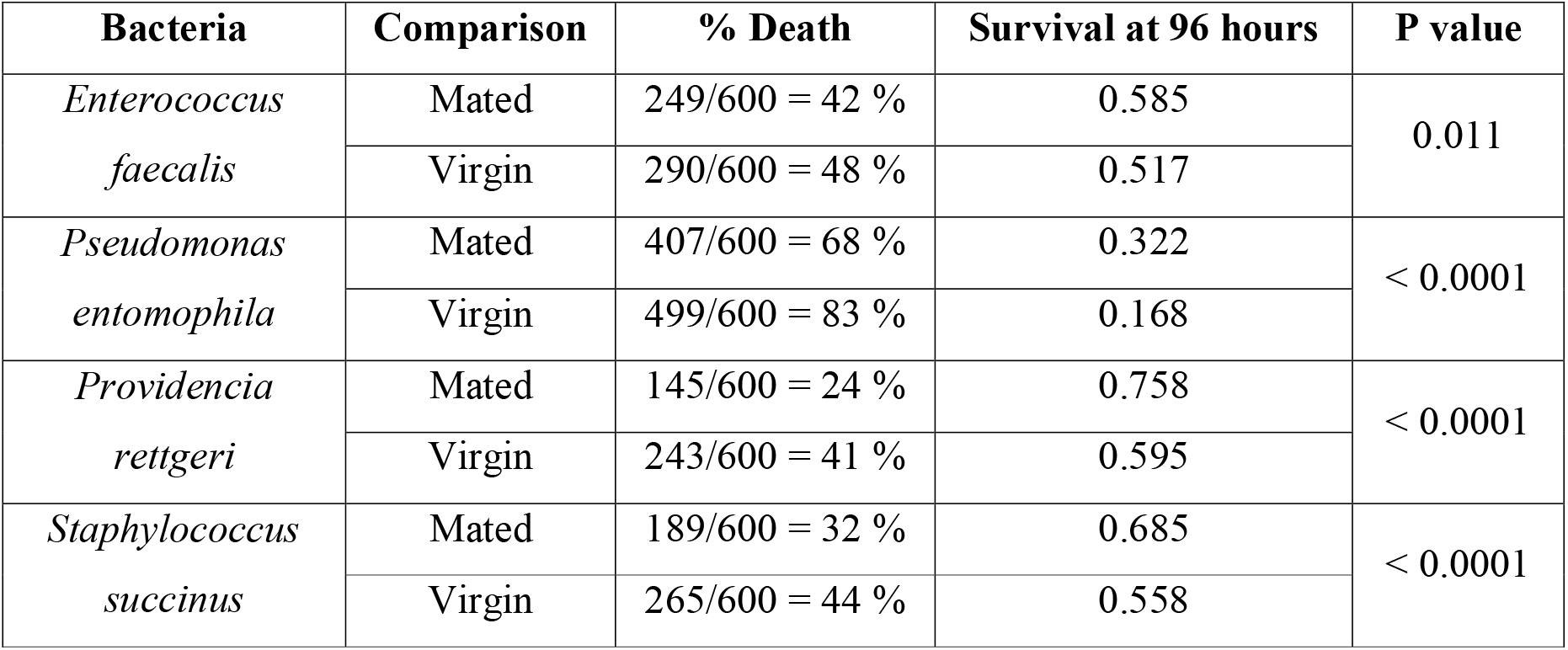
Kaplan-Meier analysis comparing mated and virgin *Drosophila* BRB flies infected with bacteria.

| Bacteria | Comparison | % Death | Survival at 96 hours | P value |
| --- | --- | --- | --- | --- |
| *Enterococcus faecalis* | Mated | 249/600 = 42 % | 0.585 | 0.011 |
|  | Virgin | 290/600 = 48 % | 0.517 |  |
| *Pseudomonas entomophila* | Mated | 407/600 = 68 % | 0.322 | < 0.0001 |
|  | Virgin | 499/600 = 83 % | 0.168 |  |
| *Providencia rettgeri* | Mated | 145/600 = 24 % | 0.758 | < 0.0001 |
|  | Virgin | 243/600 = 41 % | 0.595 |  |
| *Staphylococcus succinus* | Mated | 189/600 = 32 % | 0.685 | < 0.0001 |
|  | Virgin | 265/600 = 44 % | 0.558 |  |

## Discussion

While immunity and reproduction impart discrete fitness benefits to an organism, theory suggests a trade-off when effects of both activities exist simultaneously (Adamo, 2004; Zuk and Stoehr, 2002). Consequently, sexually active individuals may opt to bear likely risk to their immunity in lieu of potential fitness benefits from the present reproductive events, at least over a short period. We tested this hypothesis by infecting mated and virgin flies of three *D. melanogaster* BRB replicates with two Gram-positive bacteria (*Enterococcus faecalis* and *Staphylococcus succinus*) and two Gram-negative bacteria (*Providencia rettgeri* and *Pseudomonas entomophila*). We ensured multiple mating for experimental flies since previous study have shown no difference in immune function between singly mated and virgin *D. melanogaster* flies (Gupta et al., 2013).

Contrary to our expectation that the mated BRB flies would be immunocompromised compared to virgins as demonstrated in previous studies (Fedorka et al., 2007; McKean and Nunney, 2001; Short and Lazzaro, 2010), we found that the mated BRB flies have significantly higher probability of surviving all four bacterial infections (see Figure 1 and Table 1). Although coalignment between reproduction and immunity occur in the females of other insects (Barribeau and Schmid-Hempel, 2017; Msaad Guerfali et al., 2018; Shoemaker et al., 2006; Valtonen et al., 2010; Worthington and Kelly, 2016), only one study has found such trend in *D. melanogaster* BRB males suffering from *Pseudomonas entomophila* infection (Gupta et al., 2013). In the same study (Gupta et al., 2013), *Staphylococcus succinus* infection did not elicit a difference in survival between mated and virgin BRB males. However, our study did observe a better immunocompetence in mated BRB flies relative to virgins even for *Staphylococcus succinus* infection. The reason for such differences in result is difficult to know but it is important to note the 150 generations gap of fruit fly evolution between the two studies. In conclusion, although disease resistance is often pathogen specific (Adamo, 2004), we found mating enhanced immunity in *D. melanogaster* to sub lethal doses of different bacteria instead of a trade-off between reproduction and immunity.

Immunocompetence is generally thought to differ between sexes due to males’ propensity to maximize current multiple mating opportunity ignoring immunity expenditure (Zuk and Stoehr, 2002). Food availability being similar to both sexes of BRB files, multiple sexual encounters in 48 hours mating period should have led female biased immunocompetence as previously demonstrated (McKean and Nunney, 2005). Our result shows that the females did not differ from males in their ability to survive bacterial infections (see Figure 2). This trend corroborates with the alternative theory that suggests a dynamic interaction between males and females with no particular advantage for either sexes (McKean and Nunney, 2005). An explanation for similarity in immunity between sexes could be absence of known immunosuppressants in invertebrates unlike vertebrates (Letitia et al., 2000). It appears that BRB flies do not have sexual dimorphism in immunity despite having sex specific mating strategy and a promiscuous mating system.

The ability to ultimately survive potential infections from natural pathogens have implications on the evolution of a species. Since all four bacteria used were natural pathogen to *D. melanogaster*, they are likely to impose selection pressure on *Drosophila* immunity. An improvement in immunity due to mating could be an adaptation resulting from natural or sexual selection or both that eventually increases an individual’s Darwinian fitness. Also, any antagonistic relation between two fitness determining traits would be more evolutionarily limiting. A cost-free resistance to immunity may also explain the lack of trade-off between immunity and reproduction in our study (Rigby et al., 2002). Although the underlying mechanisms for improved bacterial resistance observed in mated BRB individuals remain unclear, we are able to show the true nature of interaction, i.e. synergy between immunity and reproduction, by using wild type flies infected with its natural pathogens.

## Acknowledgement

We specially thank Manas Samant and Aabeer Kumar Basu for their assistance during this study. Nitin Bansal and Tejashwini Hegde acknowledge KVYP fellowship, Biswajit Shit acknowledge INSPIRE fellowship and Aparajita acknowledge UGC-NET scholarship for support throughout the study. To the other members of Evolutionary Biology Laboratory at Indian Institute of Science Education and Research Mohali, we shower tribute for their omnipresent generosity.

